# High-throughput analysis of ANRIL circRNA isoforms in human pancreatic islets

**DOI:** 10.1101/2022.01.03.474854

**Authors:** Hannah J. MacMillan, Yahui Kong, Ezequiel Calvo-Roitberg, Laura C. Alonso, Athma A. Pai

**Author notes:** The authors wish it to be known that, in their opinion, the first 2 authors should be regarded as joint First Authors. Yahui Kong, Curia Global, Inc., Hopkinton, MA, 01748, USA.

## Abstract

The <u>a</u>ntisense <u>n</u>on-coding <u>R</u>NA in the <u>I</u>NK <u>l</u>ocus (ANRIL) is a hotspot for genetic variants associated with cardiometabolic disease. We recently found increased ANRIL abundance in human pancreatic islets from donors with certain Type II Diabetes (T2D) risk-SNPs, including a T2D risk-SNP located within ANRIL exon 2 associated with beta cell proliferation. Recent studies have found that expression of circular species of ANRIL is linked to the regulation of cardiovascular phenotypes. Less is known about how the abundance of circular ANRIL may influence T2D phenotypes. Herein, we sequence circular RNA in pancreatic islets to characterize circular isoforms of ANRIL. We identify highly expressed circular ANRIL isoforms whose expression is correlated across dozens of individuals and characterize ANRIL splice sites that are commonly involved in back-splicing. We find that samples with the T2D risk allele in ANRIL exon 2 had higher ratios of circular to linear ANRIL compared to protective-allele carriers, and that higher circular:linear ANRIL was associated with decreased beta cell proliferation. Our study points to a combined involvement of both linear and circular ANRIL species in T2D phenotypes and opens the door for future studies of the molecular mechanisms by which ANRIL impacts cellular function in pancreatic islets.

## INTRODUCTION

Type 2 diabetes (T2D) is a worldwide problem of increasing social and economic significance. Although obesity and insulin resistance are important contributors, an essential common element required for T2D progression is failure of pancreatic islets to produce sufficient circulating insulin to meet demand (1). Large scale efforts have identified >400 genomic regions that are associated with T2D risk in human populations (2); however, identifying the underlying molecular mechanisms has been slower than hoped.

Polymorphisms at the *CDKN2A/B* gene locus at chromosome 9p21 are prominently associated with diabetes risk in disparate human populations and across a range of T2D-related syndromes (3). Although the gene products at *CDKN2A/B* play roles in many cell types, strong evidence links *CDKN2A/B* T2D risk-polymorphisms specifically to impaired insulin secretory function in humans, pointing to impact on pancreatic islets as the causative mechanism by which this gene locus drives diabetes risk (4–10).

The *CDKN2A/B* locus contains two protein-coding genes: *CDKN2A*, which encodes p14^ARF^ and p16^INK4A^, and *CDKN2B* which encodes p15^INK4B^, and a long noncoding RNA called *CDKN2B-AS* or *ANRIL* (11). Although p14^ARF^, p16^INK4A^ and p15^INK4B^ play multiple biological roles in mouse and human β-cells (11) and were widely assumed to mediate the T2D risk transmitted by risk-SNPs at this locus, a study we performed analyzing *CDKN2A/B* risk-SNP impact on human islets showed a stronger association with *ANRIL* than with the protein-coding genes (12). Specifically, in 95 unique nondiabetic human islet samples, two T2D risk alleles at *CDKN2A/B* were differentially associated with *ANRIL* abundance in an age-dependent manner (12). Intriguingly, carriers of a T2D risk-SNP located within an exon of *ANRIL* had a reduced β-cell proliferation index than carriers of the protective allele (12). These findings outline an urgent need to better understand the biology of *ANRIL* in pancreatic islets.

*ANRIL* is a low-abundance but widely expressed lncRNA that is implicated in multiple disease states and diverse cell types (13–15). Containing at least 22 exons (16), many *ANRIL* isoforms have been described, including circular forms (13, 17–21), and are distributed across both nuclear and cytoplasmic compartments (13, 18, 22–27). ANRIL expression has previously been associated with cell growth, proliferation, and apoptosis phenotypes in cancer, senescence and cardiovascular disease cellular models (14, 28, 29). Intriguingly, increasing evidence suggests that there may be cross-talk or divergent functionality between linear and circular isoforms of ANRIL. Linear ANRIL (linANRIL) is thought to bind to PRC1/2 and act as a molecular scaffold guiding epigenetic complexes to promote cell adhesion, proliferation, and apoptosis (21). However, circular ANRIL (circANRIL) instead appears to compete with ribosomal RNA (rRNA) for binding with PES1 - a component of the PeBoW complex that plays a role in cell proliferation via pre-rRNA processing - resulting in impaired rRNA maturation and inhibition of proliferation (20, 21). Consequently, the relative abundance of linANRIL and circANRIL may regulate proliferative or apoptotic phenotypes associated with metabolic disease and has been implicated as such in models of atherosclerosis risk (17, 20), cancer proliferation (18), and epithelial cell response to stimulus or injury (30).

Despite the increasing recognition that circular forms of ANRIL may play a role in disease biology, little is known about the mechanisms that underlie circANRIL expression. Circular RNAs are produced by non-canonical splicing between a downstream 5’ splice site with an upstream 3’ splice site, in a mechanism called back-splicing. Since these molecules have no 5’ cap or 3’ polyA tail, they are not believed to be translated and cannot be targeted by exonucleolytic degradation. Thus, circANRIL isoforms are thought to have greater stability than linANRIL species (17, 20, 30) and, consistently, have been seen at greater levels than linANRIL in immune cells (17, 20). The handful of studies that have characterized ANRIL exons involved in back-splicing events have not identified any cis-elements that may promote back-splicing. However, most of these studies lacked the ability to perform a fully unbiased identification of circANRIL isoforms and their relative expression levels, potentially obscuring the full picture of exons and genomic elements that may contribute to ANRIL circularization. Here, we use high-throughput sequencing to identify and quantify circular ANRIL isoforms in human pancreatic islets.

## MATERIAL AND METHODS

### Human pancreatic islet preparations

Human pancreatic islets were obtained from the Integrated Islet Distribution Program (IIDP) at the City of Hope, supported by the National Institute of Diabetes and Digestive and Kidney Diseases (NIDDK), National Institutes of Health, or from a collaborative group headed at Vanderbilt University (31). Human islet studies were determined by the University of Massachusetts Institutional Review Board to not qualify for institutional review board review or exemption because they do not involve the use of human subjects.

De-identified islet samples from 112 subjects without diabetes and 10 subjects with T2D were live shipped in Prodo islet transport media. Human islets were cultured overnight in islet culture medium for recovery from isolation and shipment, then were handpicked and flash frozen at -80°C as previously described (12). DNA and RNA from the frozen islets were extracted using the Norgen RNA/DNA/Protein Purification Kit (Norgen Biotek Corp. Thorold, Ontario, Canada).

### RNA sequencing from islet preparations

RNA from 5 frozen islet donor preps acquired via the IIDP was isolated using TRIzol (15596018, Thermo Fisher Scientific). All RNA integrity numbers (RIN), as measured on a 4150 Tapestation System using a High Sensitivity RNA ScreenTape Assay kit (5067-5579/5580, Agilent Technologies), were greater than 8. Ribosomal RNA depletion was performed with the Illumina TruSeq Ribo-Zero Gold rRNA Removal kit (MRZH116, Illumina). Following the method in (32), rRNA depleted islet RNA was digested using RNase R (RNR07250, Lucigen) or mock digestion (protocol without RNase R) followed by clean-up with the Qiagen RNeasy Mini kit (Q74106, Qiagen). RNA-seq libraries were prepared with the TruSeq Stranded Illumina Total RNA Preparation Kit (20020599, Illumina). Paired-end 2×150nt sequencing of libraries was performed in-house on the Illumina NextSeq 550 sequencer using a NextSeq 500/550 300 cycle High Output Kit v2.5 (20024908, Illumina).

### Analysis of RNA-seq data and gene expression

Quality control for the RNA-seq data was performed using FastQC, files were filtered for low quality reads, and adapters were trimmed using trimmomatic/0.3.2 (33). Data were mapped to the hg38 reference genome (16) using STAR/2.7.0e (34), guided by gene annotations from ENSEMBL hg38.v95. Data from Haque *et al*. (35) were downloaded from the NCBI BioProject Database (accession number PRJNA607015) and mapped as described above. Linear RNA gene expression levels were measured using TPMs generated by kallisto/0.4.0 (36).

### Identifying and quantifying circular RNAs from RNA-seq data

Back-splice junctions (BSJs) were first identified in the RNA-seq data with the CIRCexplorer2 (37) characterization pipeline. The CIRCexplorer2 Parse module was run with default parameters using chimeric alignment output files created by using the --chimOutType flag when running STAR to create back-splice junction files. Sashimi plots in Figure 1A were generated using the following data: (1) normalized base-specific read counts per strand, generated with igvtools count, (2) linear splice junction reads reported in the sj.out.tab file from the STAR aligner, and (3) back-spliced junctions reported by CIRCexplorer2. To quantify circRNA abundance, we use the RNase R effect correction pipeline in the CIRIquant package (38). This package provides the ability to perform a direct comparison between linear and circular RNA species and accounts for both RNase R and untreated samples to calculate an adjusted BSJ count per locus. High-confidence BSJs were defined as those present in more than one individual and/or supported by more than one junction read.

**Figure 1.**
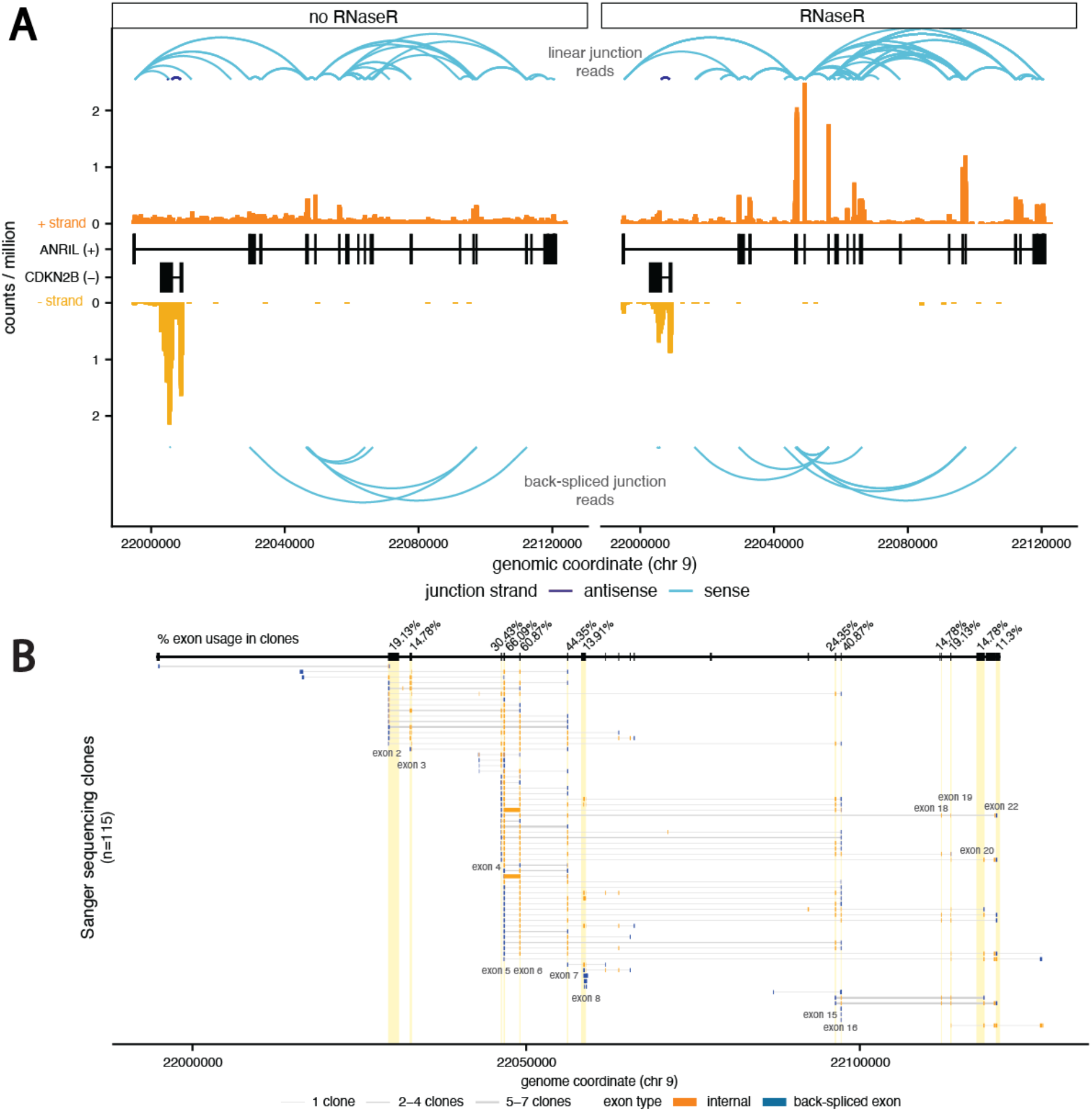
Transcription of ANRIL locus results in diverse circular RNAs. **(A)** RNA-seq data from the CDKN2B/CDKN2B-AS locus from control (*left*) and RNaseR treated (*right*) RNA. Exonic read coverage is in orange shades and splice junction reads are in blue, differentiated by linear (*top*, from STAR alignment) and backspliced (*bottom*, from CIRCexplorer2 alignment) junction reads. All read counts are normalized by the total number of reads in the libraries **(B)** Visualization of individual fragments identified by Sanger sequencing, with each row representing one isoform and the thickness of the lines representing the number of corresponding sequencing clones. Exons involved in back-splicing are represented in blue.

The ANRIL meta-isoform was created by merging all overlapping isoform annotation coordinates for linear CDKN2B-AS1 isoforms in ENSEMBL v95. Adjusted BSJ reads from CIRIquant were counted and attributed to annotated ANRIL exons based on splice site usage. HTseq/0.1.0 (39) was used to count exonic coverage across RNase R and untreated samples.

To quantify the total expression of circRNAs within each sample, we calculated a junctions per million mapped reads (JPM) metric, based on the principle that each BSJ read could only be derived from a single circRNA molecule: JPM = (# BSJ reads)/(# mapped reads * 10^6^).

### Sequencing validation of ANRIL circular RNAs

The human β cell line, EndoC-βH1, was cultured as previously described and passaged every 7 days (40). Cells were grown in culture vessels coated with 2 µg/ml Fibronectin (F1141, Sigma-Aldrich) and 1% extracellular matrix (ECM) (E1270, Sigma-Aldrich) mixed in high glucose DMEM (11965-092, Life Technology) and cultured in low glucose DMEM containing 5.5 mM glucose (11885-084, Life Technology)), 2% bovine serum albumin (BSA) (BAH66, Equitech), 10 mM nicotinamide (481907, VWR), 100 U/ml penicillin, 100 µg/ml streptomycin (P/S) (P4333, Sigma-Aldrich), 50 μM β-2-mercaptoethanol (M3148, Sigma-Aldrich), 5.5 μg/ml transferrin (T8158, Sigma-Aldrich)and 6.7 ng/ml sodium selenite (S1382, Sigma-Aldrich).

Total RNA was isolated from EndoC cells using the Norgen RNA/DNA/Protein Purification Kit (Norgen Biotek Corp. Thorold, Ontario, Canada) and subjected to reverse transcription using the SuperScript IV VILO MasterMix kit (Thermo Fisher Scientific) according to the manufacturer’s protocols. Outward facing primers were used against ANRIL exons 2, 3, 4, 6, 7, 8, 16 and 19 with forward primer against the 3′ end of each exon and reverse primer against 5′ end of the same exon (Table S3). PCR reactions were performed following the cycling conditions: 95°C for 5 min, 35× (95°C for 30 s, 59°C for 30 s, and 68°C for 1 min), and 68°C for 5 min. PCR products were cloned using the TOPO TA Cloning Kit (Life Technologies). The positive clones were selected by digesting the purified clones with XhoI and HindIII restriction enzymes and sequenced using M13 Forward (−21) and M13 Reverse primers performed in two directions. The sequencing results from the clones were aligned to each exon sequence of ANRIL to identify the circular ANRIL junctions and isoforms.

### qRT-PCR quantification of linear and circular RNA

Total RNA was reverse transcribed using a SuperScript IV VILO MasterMix kit (Thermo Fisher Scientific). For relative expression of circANRIL junctions in EndoC cells and human islet samples by qPCR, Taqman primer and probe sets for the detection of ANRIL junctions 7–5, 10-5, 16–5, and 16-4 were designed. Together with two commercial TaqMan ANRIL expression assays ANRIL (Exon5-6), Hs04259476_m1; and ANRIL (Exon1-2), Hs01390879_m1, as well as those for CDKN2B p15INK4b, Hs00793225_m1; CDKN2A p16INK4a, Hs02902543_mH; CDKN2A ARF, Hs99999189_m1, the expression levels of target genes were quantitatively assessed in duplicate by normalizing to the level of endogenous reference (ACTB, Hs01060665_g1; and GAPDH, Hs02758991_g1) Transcript expression levels were presented as log2-transformed expression (ΔCt).

#### RNase R treatment for qPCR

Equal amounts of RNA (2µg) were incubated with or without 20 U of Rnase R (RNR07250, Epicentre Biotechnologies) and 40 U of RiboLock RNase inhibitor (EO0381,Thermo Fisher Scientific) in a 20µL reaction volume for 30 minutes at 37°C. The resulting RNA was purified as described above and quantified. Equal volumes of RNA were then subjected to reverse transcription using the SuperScript IV VILO MasterMix kit (Thermo Fisher Scientific, #11756050) with a mixture of random hexamer and oligo dT primers. qPCR was performed to quantify the transcripts using Taqman primers/probes for ANRIL exon1-2, ANRIL exon5-6, ANRIL back-spliced junctions (7-5, 10-5, 16-4 and 16-5). Actin and GAPDH were used as controls. The mRNA depletion was examined by normalizing the relative expression of transcripts with RNase R treatment to the untreated control.

#### Isolation of cytoplasmic and nuclear RNA

EndoC cells were collected for subcellular fractionation and total RNA isolation. Cytoplasmic and nuclear RNA were isolated with the Cytoplasmic & Nuclear RNA Purification Kit (Norgen, Belmont, CA, USA) following the manufacturer’s manual. Briefly, EndoC cells were harvested and incubated with a lysis buffer for 5 min on ice. Then, the cells were centrifuged at maximum speed for 10 min at 4°C, the supernatant was kept for assessing the cytoplasmic RNA, and the pellet was used for nuclear RNA extraction. Next the RNA was reversely transcribed to cDNA according to the instructions of SuperScript IV VILO MasterMix kit (Thermo Fisher Scientific). The expression levels of ANRIL (Exon5-6), Hs04259476_m1; ANRIL (Exon1-2), Hs01390879_m1, circular ANRIL (7-5, 10-5, 16-4 and 16-5) in the whole cells, nuclei and cytoplasm were examined by qRT-PCR. Glyceraldehyde 3-phosphate dehydrogenase (GAPDH) and Nuclear Enriched Abundant Transcript 1 (NEAT1) were detected as fractionation indicators. The primers for GAPDH RNA were 5’-CTCCTCCTGTTCGACAGTCA-3’ (sense) and 5’-GTTGACTCCGACCTTCACCT-3’ (antisense). The primers for NEAT1 RNA were 5’-GTGGCTGTTGGAGTCGGTAT-3’ (sense) and 5’-TAACAAACCACGGTCCATGA-3’ (antisense).

### Sequence Characteristics for ANRIL exons and introns

#### Splice site scores

The strength of splice sites for ANRIL exons were calculated using a maximum entropy model as implemented in maxEntScan (41) using 9nt around the 5’ splice site (−3:+6) and 23nt around the 3’ splice site (−20:+3).

#### Repeat regions

Repeat annotations for all repeat classes were downloaded from RepeatMasker (www.repeatmasker.org (42)) and overlapped with ANRIL meta-exon and intervening intronic coordinates to look for enrichments within regions involved in or proximal to back-splicing events.

#### Sequence complementarity

ANRIL exon and intron sequences were extracted based on the ANRIL meta-isoform coordinates and inverted complementary sequences were created for each region. All pairwise combinations of exon-exon or intron-intron (for upstream or downstream flanking introns, independently) were evaluated for sequence complementarity using blastn (parameters -word_size 7 - gapopen 5 -gapextend 2 -penalty -3 -reward 2) similar to described previously (43). Pairs of regions seen to interact in BSJs were compared to an equal number of randomly selected non-occurring pairings.

### Motif Analyses

ANRIL exon and intron sequences were extracted based on the ANRIL meta-isoform coordinates. For exons, full exonic sequences were used, but introns were split into two equal regions proximal or distal to the 3’ or 5’ splice site involved in the back-splicing reaction and proximal regions were only compared to similarly defined proximal regions within the splice site type. For example, intronic regions proximal to BSJ-involved 3’ splice sites were compared to intronic regions proximal to 3’ splice sites not involved or minimally involved in BSJs. Motif analyses were conducted using STREME (44) to identify enriched sequence motifs in BSJ-involved exons or proximal intron regions relative to exons or intronic regions with no or low evidence for back-splicing.

To perform similar motif analyses across the full set of circRNAs expressed in islets, we first considered genes with a TPM higher than 5 measured by kallisto/0.4.0 and identified expressed exons as those with junction read counts in the bottom 10% of the distribution of junction counts as reported by STAR/2.7.0e. Expressed exons that matched BSJ coordinates from CIRIquant were probed for enriched sequence motifs using STREME, with expressed exons not matching BSJ coordinates used as a background set. For intronic motif enrichments, we used sequences 100bp upstream of 5’ splice sites and 100bp downstream of 3’ splice sites. Enriched motif sequences for all of the above comparisons were compared with the RNA Binding Protein Database (RBPDB; http://rbpdb.ccbr.utoronto.ca/) to identify candidate RNA binding proteins that bind to enriched motifs.

### miRNA analyses

To define full length circANRIL isoforms, we considered as circular any exon present in at least 10% of clones sequenced using Sanger sequencing. We used miRanda 3.3a (45) to predict miRNA target sites in ANRIL exons, only considering miRNAs with a mean expression in islet cells greater than 5 reads per million based on three datasets downloaded from miRmine (https://guanfiles.dcmb.med.umich.edu/mirmine/index.html): SRX290576,SRX290601,SRX290617). Finally, we normalized the number of miRNA sites in each exon category (linear or circular) by the total length of the exons.

### Cellular phenotype assays

#### Proliferation assays

Human islets from a subset of donors (n=45) were cultured overnight in islet culture medium, trypsinized to single cells using and plated on uncoated glass coverslips (Fisherbrand) as previously described (12). Dispersed cells were cultured in islet culture medium containing either 5 mmol/L or 15 mmol/L glucose and 20 μg/mL bromodeoxyuridine (Sigma) for 96 hours. After culture, the islet cells were fixed for 10 min in 4% paraformaldehyde (Sigma). Fixed cells were unmasked in 1N HCl for 20 min at 37°C, blocked for 2 hours in goat serum–based block with 0.1% Tween 20. Immunofluorescence staining was performed for insulin (ab7842, Abcam, or A056401-2; Dako), and BrdU (ab6326; Abcam) antibodies, and DAPI (Sigma) as previously described (12). The percent of insulin-staining cells that were also BrdU labeled, was quantified on blinded images to calculate β–cell proliferation. Data were expressed as the proliferation index, calculated as the ratio of %BrdU+ β-cells in 15 mmol/L glucose divided by the %BrdU+ β-cells in 5 mmol/L glucose.

#### Insulin Secretion Assays

A subset of islet samples (*n* = 83) were tested for glucose stimulated insulin secretion at the time of islet isolation by each individual isolation center; these data were downloaded from IIDP website.

#### Genotyping

Genotyping for four CDKN2A/B SNPs (rs564398 [C/T], rs10811661 [C/T], rs2383208 [G/A], and rs10757283 [C/T]) were performed in the DNAs of human islets using commercial (C_2618017_10, C_31288917_10, C_15789011_10, and C_31288916_10) TaqMan SNP genotyping assays (Thermo Fisher Scientific, Waltham, MA) as previously described (12).

### Gene Ontology Analyses

Gene Ontology analyses were conducted by associating each BSJ with an individual ensembl gene and identifying the following subsets: top 10% and bottom 10% of expressed circRNAs based on JPM values and top 10% and bottom 10% of circular/linear ratios based on JPM/TPM ratios. Background sets were all expressed genes and all expressed genes with circRNAs, respectively. Gene ontology enrichment analyses were performed in an iterative fashion with a custom script to avoid significant gene ontology terms with overlapping gene sets, as described before (46). P-values were computed using a Fisher-exact test and then corrected using a Benjamini-Hochberg multiple test correction.

## RESULTS

Previous studies in immune, cancer, and epithelial cells have identified a wide range of circANRIL isoforms that are expressed in different conditions (17, 18, 20, 30). However, such an analysis has not been conducted in pancreatic islets, despite evidence that circular and linear ANRIL expression may be implicated in diabetes phenotypes. The aim of our study is to systematically characterize circANRIL isoforms and their regulation in non-diabetic human islet tissue, with the goal of defining the circANRIL landscape in this critical metabolic tissue.

### Identifying circular isoforms of ANRIL in pancreatic islets

We first set out to identify pancreatic islet circANRIL isoforms in an unbiased fashion. To do so, we obtained five frozen islet preparations from the Integrated Islet Distribution Program (Methods; **Table S1**) and conducted RNA-seq after digestion with RNase R (which enriches for circRNA by degrading linear RNA molecules), paired with mock digested samples to obtain total RNA abundance for each sample. To identify back-spliced exon junctions indicative of circularization, we used the circRNA analysis package Circexplorer2 (37) after linear read alignment with STAR (34). Supporting efficacy of this approach, the RNase R treated samples had over 4.5x more back-spliced junctions (BSJ) reads than the untreated samples (0.65% vs 0.14% of all reads, respectively; **Figure S1A**). RNA-seq data across different islet samples were highly consistent, with an average gene expression correlation greater than 0.95 within RNase R treated samples and 0.94 within untreated samples (Pearson correlations, **Figure S1B**). Our data correlated well with publicly available pooled human islet RNA-seq from Haque *et al*. (35), providing additional confidence to our sample preparation. Thus, we decided to include the Haque *et. al*. data in our downstream analyses to provide more statistical power for circANRIL isoform identification. To identify circANRIL isoforms independent of known linear isoforms, which contain different subsets and combinations of exons, we collapsed all annotated linear exons into a meta-isoform and re-numbered exons based on this meta-isoform for reference in all downstream analyses (**Figure S1C**).

Upon RNase R treatment, we observed an enrichment of circRNA at the ANRIL locus (**Figure 1A**). Though there were almost twice as many linear junction reads in the untreated samples, there was a 2.65 fold increase in BSJs at the ANRIL locus in the RNase R treated samples (**Table S2**). Interestingly, there were also more linear junction pairs of splice sites in the RNase R samples, perhaps indicating greater alternative splicing in circRNA molecules. Reassuringly, read coverage for the antisense CDKN2B mRNA was lower in the RNase R treated samples and no BSJ reads mapped to this locus. A number of ANRIL exons had greater read coverage in the RNase R samples, suggesting that ANRIL circularization might preferentially involve specific ANRIL exons. To quantify circRNA abundance we used the CIRIquant package (38), which performs a direct comparison between linear and circular RNA species and accounts for both RNase R and untreated samples to calculate an adjusted BSJ count per locus (**Table S3**). After removing BSJs that occurred in only one individual or were only supported by one read, we identified 17 high confidence circANRIL BSJs that used splice sites corresponding to 11 different exons and three intragenic regions (**Figure S1D**). To validate our high-throughput sequencing data, we designed divergent primers in 8 ANRIL exons and conducted PCR, cloning, and Sanger sequencing for 115 clones in the immortalized endoC-βH1 cell line (**Figure 1B**). While the lower throughput sequencing identified 35 back-splicing events involving a greater number of exons, 59% of our RNAseq-based BSJs were validated, including all of the BSJs with the greatest read coverage (**Figure S1E**). Interestingly, Sanger sequencing clones also identified extensive ANRIL cryptic splice site and exon usage, with a number of intronic sites involved in back-splicing and intronic sequences spliced into circANRIL isoforms (**Table S4**).

### Specific circANRIL isoforms are highly abundant

Next, we capitalized on the long-read Sanger sequencing data to gain insight into the internal exon structure of circANRIL isoforms. Since Sanger sequencing data is biased by the number of clones sequenced from each primer set (which are placed in exons that are likely not equally represented across circANRIL isoforms), these data do not provide quantitative information about isoform expression. Nevertheless, we observed a striking concordance: exons that were present in the majority of Sanger clones were also overrepresented in the RNase R RNA-seq data, measured by the fold change in exonic coverage between RNase R and untreated samples (**Figure 2A**) These overrepresented exons also tended to fall within the linear boundaries of exons that were most frequently used in back-splicing reactions, including exons 2, 4 and 5 as 3’ acceptors and exons 7, 10, and 16 as 5’ donors in the back-splicing reactions.

**Figure 2.**
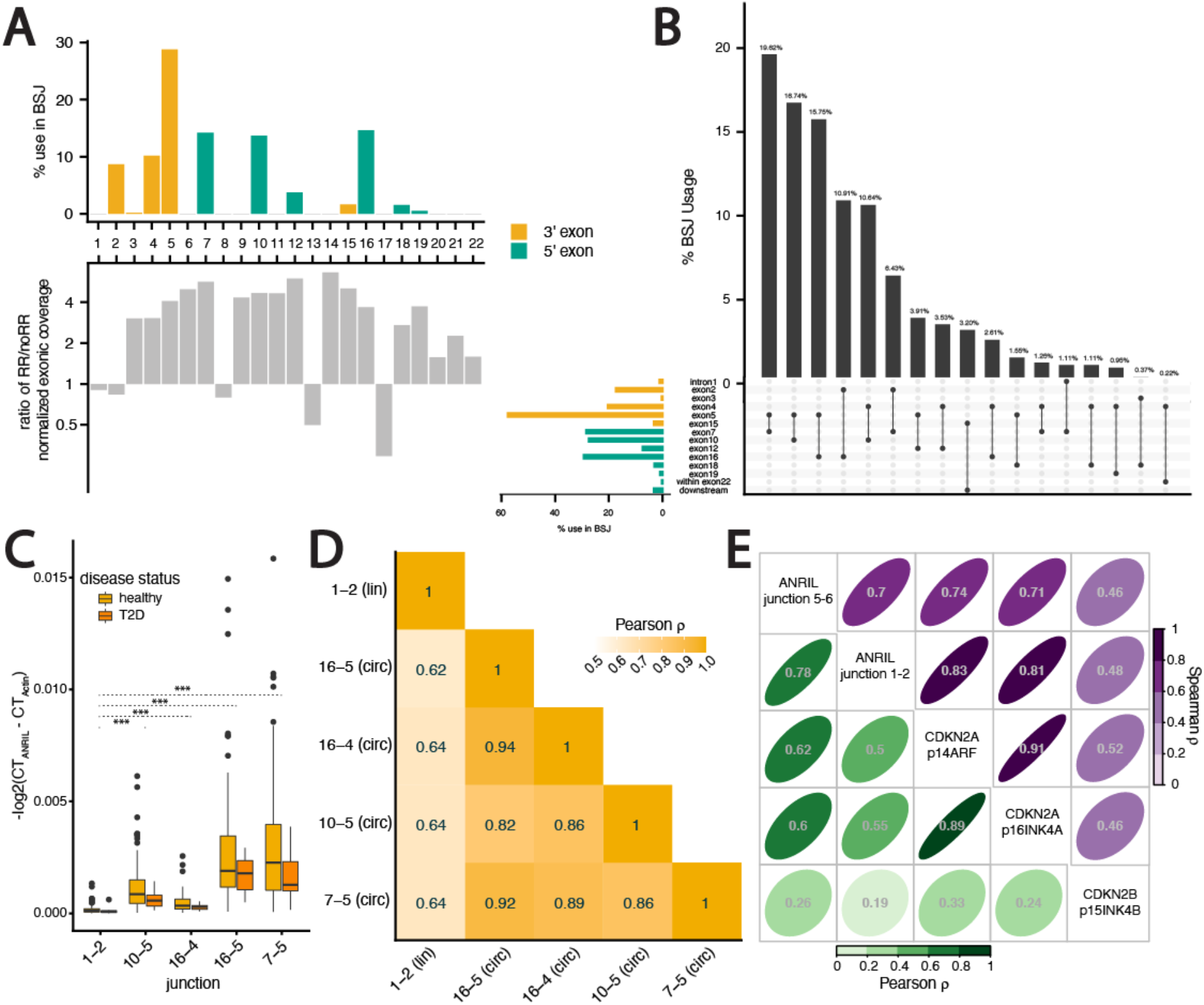
Specific circANRIL isoforms are highly abundant. **(A)** Exon-specific involvement in back-spliced junctions (*top*) and fold-change in expression between RNaseR treated and control samples (*bottom*, normalized for library depth). **(B)** Frequency of splice site pairing in back-spliced junctions, with proportional usage (*top*) for each BSJ (*bottom*), with proportional usage of individual exons on the left. **(C)** Pairwise Pearson correlations between linear (junction 1-2) and circular ANRIL expression across 122 islet samples. **(D)** RT-PCR quantification of linear and circular isoforms across 122 islet preparations. **(E)** Pairwise Pearson correlations (*bottom left*) and Spearman rank correlations (*top right*) between linear ANRIL (junction 1-2), circular ANRIL (junction 5-6), CDKN2A isoforms, and CDKN2B abundance across 122 islet preparations.

The overrepresentation of specific ANRIL exons in BSJs and internal circRNA structure suggests preferential expression of particular circANRIL isoforms. Indeed, we find that several BSJs occurred at higher frequency **(Figure 2B)**, with the exon7-exon5 pairing alone contributing almost 20% of BSJ reads, 4 other pairings contributing more than 10% of BSJ reads, and exon5 acting as a 3’ acceptor site in more than half of the BSJ reads. Notably, three isoforms previously seen to be predominantly expressed across immune and cancer cells types (17, 18, 20), 7-5, 10-5 and 16-5, were the three highest expressed isoforms in islets. To validate relative expression of high to moderately expressed circANRIL isoforms, we used qRT-PCR to quantify the enrichment of 4 BSJ junctions (7-5, 10-5, 16-4, and 16-5), 2 linear junctions (1-2 and 5-6) and two predominantly linear housekeeping genes following RNase R treatment in endoC-βH1 cells. The relative abundance of BSJ-containing RNA remained high after RNase R treatment, while the housekeeping genes and linear 1-2 junction were expressed at significantly lower levels (all Mann-Whitney U test p-values < 0.001, **Figure S2A)**. Interestingly, relative expression of the linear 5-6 junction was similarly abundant to the BSJ containing RNA, indicating that exons 5 and 6 together may be primarily present in circRNA and serve as a good proxy for circRNA expression. This is consistent with the notable increase in reads for exons 5 and 6 after RNase R treatment (**Figure 1A**) and presence of these exons in greater than 65% of Sanger sequenced clones (**Figure 1B**).

The skewed frequency of splice site pairing suggested that ANRIL back-splicing may be a regulated event. Since our RNA-seq data were not powered to evaluate individual-specific expression levels across isoforms, we turned to a larger cohort of 95 islet preparations in which we had previously characterized linANRIL expression (12) and added 27 samples for a total of 122 samples (Methods; **Table S5**). With this larger set of samples, we used qRT-PCR to measure expression of linear ANRIL, using an assay for exon 1-2 found predominantly in linANRIL, and specific circANRIL isoforms identified with the RNA-seq data (Methods). First, we confirmed that circANRIL isoforms are expressed at significantly higher levels than linear ANRIL on average across these islet samples (all T-test p-values < 10^−8^**; Figure 2C)** (17, 20, 47). We see no significant differences between ANRIL isoform abundances between the non-diabetic and 10 diabetic samples in our dataset. We found that circANRIL isoforms correlated better with other circANRIL isoforms (all Pearson R > 0.8) than with linear ANRIL (average Pearson R = 0.64; **Figure 2D**), suggesting independent regulation of back-splicing across individuals after transcription at the ANRIL. To understand the regulation of circANRIL, we examined the Spearman rank correlations between circANRIL expression (measured as junctions per million, JPM; Methods), and the expression of other RNAs transcribed at the locus. Across the 5 islet samples on which we performed RNA-seq, we found a lower correlation between linear and circANRIL than with other genes transcribed at the locus **(Figure S2B)**, including the antisense mRNA CDKN2B and the upstream divergently transcribed mRNA CDKN2A. To gain more power, we used qRT-PCR measurements in the larger cohort of 112 islet samples and again found that linear and circANRIL had a lower rank correlation with each other than other genes transcribed at the locus (Spearman R = 0.7; **Figure 2E**). Consistent with our previous results (12), the expression of both linear and circANRIL are more highly correlated to the expression of CDKN2A isoforms than CDKN2B, suggesting that ANRIL and CDKN2A transcription might be co-regulated in islets by the same promoter or enhancer elements. Notably, while the Spearman rank correlations are high, the lower Pearson correlations suggest differences in transcriptional output from the ANRIL and CDKN2A loci and again point to the regulation of circANRIL expression across individuals. Finally, the low correlation between ANRIL and CDKN2B expression levels does not support a model in which ANRIL RNA directly regulates CDKN2B levels through complementary binding interactions.

### Local sequence elements may mediate circANRIL production

We observe that back-splicing is more likely to occur at a small number of splice sites at the ANRIL locus. Furthermore, the expression levels of these circular isoforms are highly correlated with each other but not with the levels of linear ANRIL. Taken together, these results suggest co- or post-transcriptional regulation of back-splicing at the ANRIL locus. To investigate potential cis-elements that regulate the formation of circRNAs, we focused on 5 features: (1) splice site strength, (2) intron length, (3) repeat regions, (4) sequence complementarity and (5) enriched motifs. For these analyses, we differentiated between circANRIL 3’ exons, which are associated with the 3’ acceptor participating in the back-splicing reaction, and 5’ exons, which are associated with the back-spliced 5’ donor (**Figure 3A**). First, we observe that 5’ splice sites of circANRIL 3’ exons are significantly weaker than those of non-BSJ and 5’ exons (bootstrapped p-value = 0.03, Methods, **Figure S3A**), suggesting that 3’ exons involved in BSJs may not be well defined by canonical exon-definition pathways and thus be more likely to have unpaired 3’ splice sites available for back-splicing with downstream exons (48).

**Figure 3.**
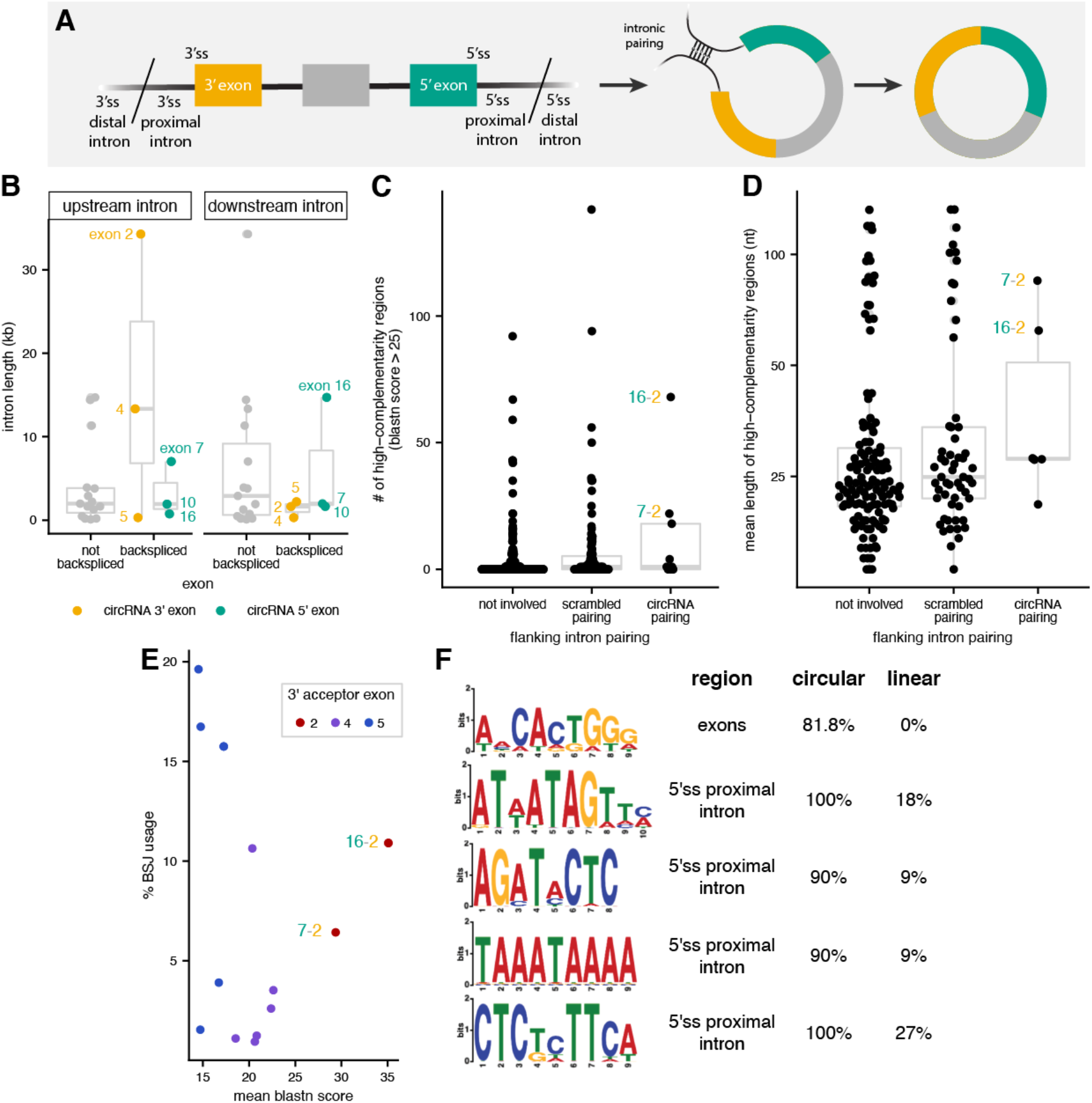
ANRIL back-splicing is associated with sequence features. **(A)** Schematic for BSJ involved 3’ and 5’ splice site (ss) proximal introns used as input for motif enrichment (*left*) and intron pairing (*right*). **(B)** Distribution of intron lengths (*y-axis*) for 3’ and 5’ back-spliced exons (*yellow* and *green*, respectively) and exons not involved in back-splicing (*grey*). **(C)** Distribution of the number of high complementarity (blastn > 25) regions (*y-axis*) within pairs of flanking introns. **(D)** Distribution of length for high complementarity regions (*y-axis*) within pairs of flanking introns. **(E)** Correlation between mean complementarity scores from blastn for high complementarity regions (*x-axis*) and %BSJ isoform usage (*y-axis*) for frequently used 3’ exons. **(F)** Sequence motifs enriched in ANRIL BSJ-involved exons or flanking introns, with percent occurrence in circular or non-circular (linear) regions indicated on the right.

Previous studies have seen that back-spliced exons tend to be flanked by longer introns (49, 50). Consistently, the introns upstream of 3’ exons 2 and 4 and downstream of exon 16 are among the longest annotated ANRIL introns (**Figure 3B**) and could potentially harbor more sequence elements that promote or regulate ANRIL back-splicing events. One feature proposed to promote back-splicing is complementarity-mediated pairing between introns flanking back-spliced exons (49, 51, 52), which brings together the upstream 3’ splice site and downstream 5’ splice site in 3D space (**Figure 3A**). Contrary to what has been observed for other circRNAs (43, 53, 54), we did not observe enrichment of known repeat regions within ANRIL introns flanking sites involved in back-splicing reactions (**Figure S3B**). Thus, we instead looked for non-repeat complementary regions in introns using blastn (Methods). We observe that the pairing between introns flanking exons 7-2 and 16-2 have among the most high-scoring complementary regions (**Figure 3C**) and the longest average length of complementary regions (**Figure 3D**) of any pairwise combinations of ANRIL introns. This suggests that the use of exon 2 as a 3’ exon in a back-splicing event may be mediated purely by sequence complementarity, consistent with the idea that exon 2 appears to mostly appear in linear ANRIL and appears in less than 10% of BSJs. Indeed, we observe that the higher-scoring 16-2 pairing is used more in BSJs than the lower-scoring 7-2 pairing (**Figure 3E**), a correlation that is not observed for the intronic pairings of other circANRIL 3’ exons.

Lastly, back-splicing may be directly regulated by splicing factors that bind to BSJ-involved exons or flanking introns (55), so we looked for enrichment of sequence motifs within BSJ involved regions relative to non-involved regions (Methods). We identified 5 motifs that are present in more than 80% of BSJ-involved exons or introns - one in exons and the other four in the intronic region proximal to BSJ-involved 5’ splice sites (**Figure 3F, S3C**). Interestingly, the motif enriched in BSJ-involved exons matches the known motif for the SFRS13A (aka SRSF10) SR-protein splicing factor, which has previously been shown to promote back-splicing in humans, drosophila, and murine models (56– 58).

### circANRIL exons have more miRNA target sites

Our results support regulated expression of ANRIL circRNA at high levels in human pancreatic islets. Thus, with our cell-type specific catalog of ANRIL isoforms, we next aimed to investigate the potential functional roles for circANRIL in islet cells. We first used the endoC cell model to ask whether ANRIL RNA was differentially localized across cellular compartments (59). Consistent with previous observations (17, 20), we find that circANRIL isoforms are predominantly cytoplasmic, with 10-fold greater abundance in the cytoplasm on average than in the nucleus (**Figure 4A**). Cytoplasmic to nuclear ratios of circANRIL are also significantly higher than linear ANRIL (all Mann-Whitney U p-values < 0.05), which is approximately equally abundant in the nucleus and cytoplasm.

**Figure 4:**
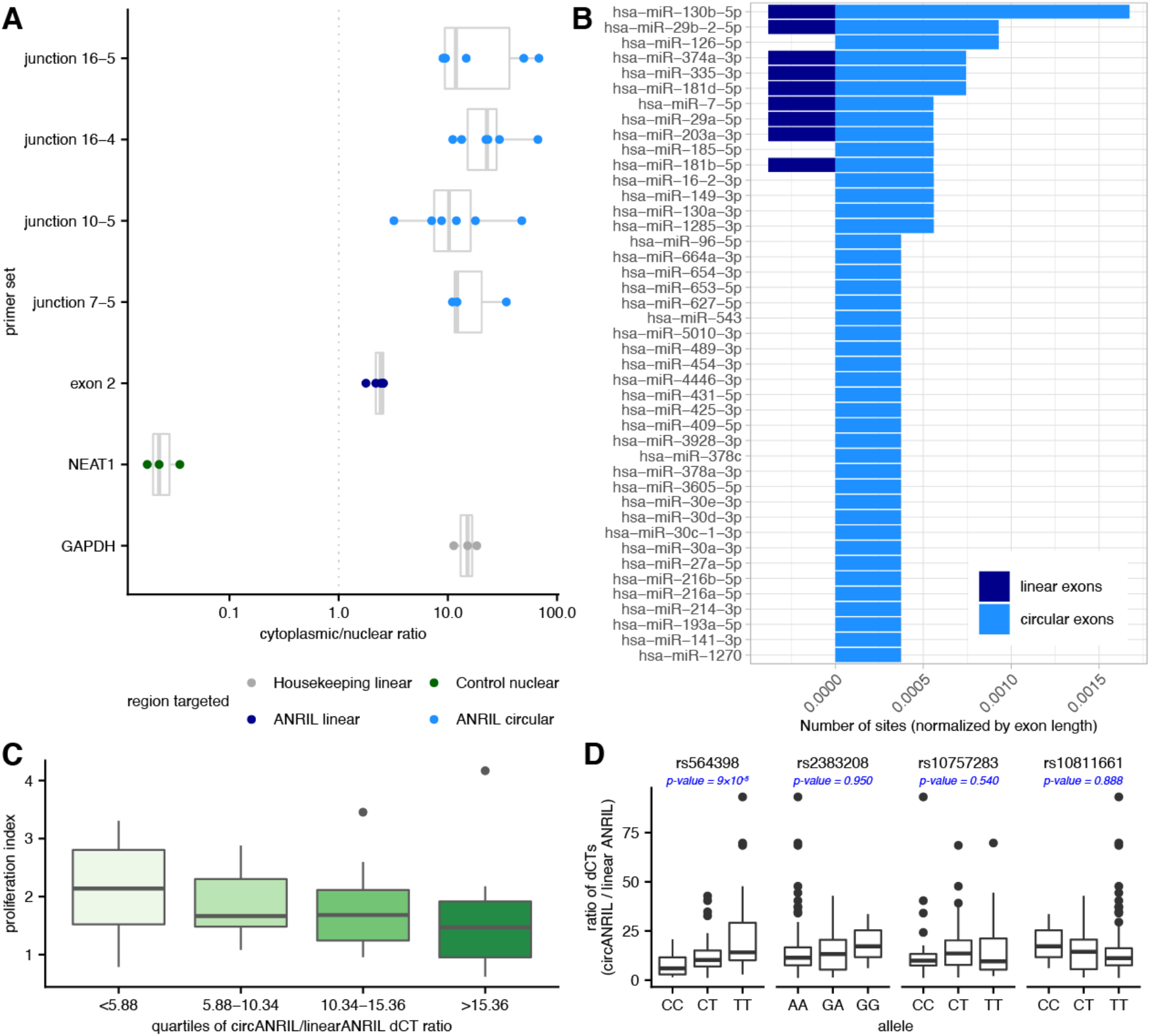
circANRIL is associated with cytoplasmic functions. **(A)** Ratio of cytoplasmic to nuclear abundance (*x-axis*) as measured by RT-PCR quantification with primer sets targeting a housekeeping gene (*grey*), a nuclear control gene (*green*), linANRIL (*light blue*) and circANRIL junctions (*dark blue*). **(B)** Number of miRNA target sites (normalized by exon length, *x-axis*) in ANRIL circular exons (*light blue*) vs ANRIL linear exons (*dark blue*) for miRNAs with more sites in circANRIL exons. **(C)** Distribution of proliferation indices (*y-axis*) for quartiles of circANRIL/linANRIL ratios (*x-axis*) as measured by RT-PCR across 45 islet preparations. **(D)** Association between genotypes and circANRIL/linANRIL ratios for 4 disease-associated SNPs at the ANRIL locus across 122 islet preparations. P-values are computed from a linear regression and Benjamini-Hochberg corrected.

circRNAs that are abundant in the cytoplasm have been previously implicated as cellular sponges acting to sequester RNA binding proteins (RBP) or miRNAs away from other regulatory targets (55, 60, 61). Since we had not found any cytoplasm RBP motifs in circANRIL exons, we turned our attention to miRNA target sites. We looked for the presence of miRNA target sites in ANRIL exons frequently used in circANRIL isoforms (both participating in BSJs or internal exons) relative to exons more frequently seen in linear ANRIL (Methods), conditioning on miRNAs that are expressed in islet cells. We find that 43 miRNAs have more target sites in circANRIL exons than linANRIL exons, while only 18 have more target sites in linANRIL exons (normalized by respective total exon lengths, **Figure 4B, S4A; Table S6**). It is important to note, however, that the total length of exons present in circANRIL is approximately double the length of exons present exclusively in linANRIL, providing greater chances for miRNA target sites to occur purely by chance. Previous experimental investigations have identified ANRIL RNA association with miR-7-5p (in periodontal ligament stem cells (62)), miR-9 (kidney epithelial cells (63)), miR-622 (brain microvascular endothelial cells (64)), miR-140-5p (lung epithelial cells (65)). miR-622 was not expressed in islets and of the remaining 3 miRNAs, only miR-7-5p was found to have more target sites in circANRIL exons. The miRNA with the highest number of sites in ANRIL exons, miR-130b-5p, has 4.38-fold more sites in circular exons than linear exons. miR-130b-5p is thought to target and downregulate several oncogenes and was seen to be downregulated itself in pancreatic cancers (66), in which ANRIL is overexpressed (67).

### Ratio of circular/linear ANRIL is associated with beta cell proliferation and diabetes risk genotype

The balance between linear and circular ANRIL expression has previously been associated with cell proliferation and apoptosis phenotypes in cancer and atherosclerosis cell models (17, 18, 20, 68, 69). Thus, we wanted to understand whether the highly abundant, cytoplasmic circANRIL isoforms in islet cells may play a role in islet biology or diabetes-relevant cellular phenotypes. We again turned to the expanded cohort of primary human islet preparations to measure two important aspects of islet function: insulin secretion and β-cell proliferation. We did not find a clear relationship between insulin secretion - measured by a glucose stimulation index (Methods) - and levels of linANRIL, circANRIL, or the relative ratio of circular to linear ANRIL abundance (circ/linANRIL; **Figure S4B**). However, we observed a clear monotonic relationship between the beta cell proliferation index and circ/linANRIL ratio. Insulin-positive cells in islet preparations expressing the lowest circ/linANRIL ratio had a significantly higher proliferation index than preparations with higher circ/linANRIL (**Figure 4C**), but no relationship was observed between proliferation and abundance of linANRIL or circANRIL isoforms alone. Previous studies have shown that over-expressing circANRIL isoforms decreased proliferation in other cell types (20, 47, 64), suggesting that the relationship between circ/linANRIL and the proliferative index in islet cells may be a consequence of increased circANRIL expression shifting the balance between the two RNA species. Since pancreatic beta cell number is an important determinant of insulin secretory capacity (70–72), and proliferation is the primary mechanism generating new beta cells in the adult pancreas (73), these observations suggest the possibility that the ratio of circANRIL to linANRIL may be one determinant of diabetes susceptibility.

Single nucleotide polymorphisms (SNPs) at the ANRIL locus associated with diabetes susceptibility have also been specifically associated with β-cell proliferation (12). Thus, we wondered whether islet circANRIL abundance was associated with genotypic diversity across individuals and if the relationship between circ/linANRIL expression and proliferation was mediated by a genetic regulatory pathway. To do so, we genotyped 4 T2D-associated SNPs at the ANRIL locus (rs564398, rs2383208, rs10757283, rs10811661) across our expanded cohort of islet preparations. Excitingly, we observed a significant association between circ/linANRIL abundance and genotype at rs564398 (**Figure 4D**), which is located within ANRIL exon2. Specifically, the T2D risk (T) allele at rs546398 was associated with higher circ/linANRIL abundance. The other 3 T2D associated SNPs, located downstream of the gene region, were not associated with circ/linANRIL. We did not see significant correlations between rs564398 genotype and the expression of either linANRIL or any circANRIL isoform independently (**Figure S4C**). The previously observed association between the homozygous protective CC genotype and higher beta cell proliferation (12) is also consistent with this genotype being associated with lower circ/lin ANRIL.

### Back-splicing is ubiquitous in pancreatic islets

To evaluate our observations about circANRIL expression within the broader context of back-splicing regulation, we looked at the global distribution of circRNA expression in human islets (**Table S7**). Consistent with previous studies (74), we found a strong negative correlation between the circular/linear ratio and linear gene expression across most genes (Spearman R = -0.6; **Figure 5A; Table S8**). However, there is a notable cluster of genes that deviate from the global trend, with moderately low linear expression but relatively high circular/linear ratios. While most of the transcripts derived from these loci are either fusion read-through transcripts, pseudogenes, HLA cluster genes, or non-coding RNAs, there are two low abundance mRNAs represented in this cluster: TAS2R14 and PHRF1. Interestingly, TAS2R14 is a bitter taste receptor family member that has been shown to stimulate GLP-1 secretion, leading to reduced β-cell apoptosis and increase in β-cell proliferation (75–78). Finally, testing for enriched sequence motifs in exons involved in back-splicing or intronic regions flanking back-spliced exons (Methods) did not reveal any motifs that overlap those enriched in ANRIL back-spliced exons (**Figure 5B; S5A**). This potentially suggests that ANRIL circularization is regulated in a locus-specific fashion rather than by global patterns. However, we did observe a number of motifs that are significantly enriched in circularized exons and respective flanking intronic regions, suggesting that there may be cis-elements that are shared between back-splicing events and regulated by common trans-factors.

**Figure 5.**
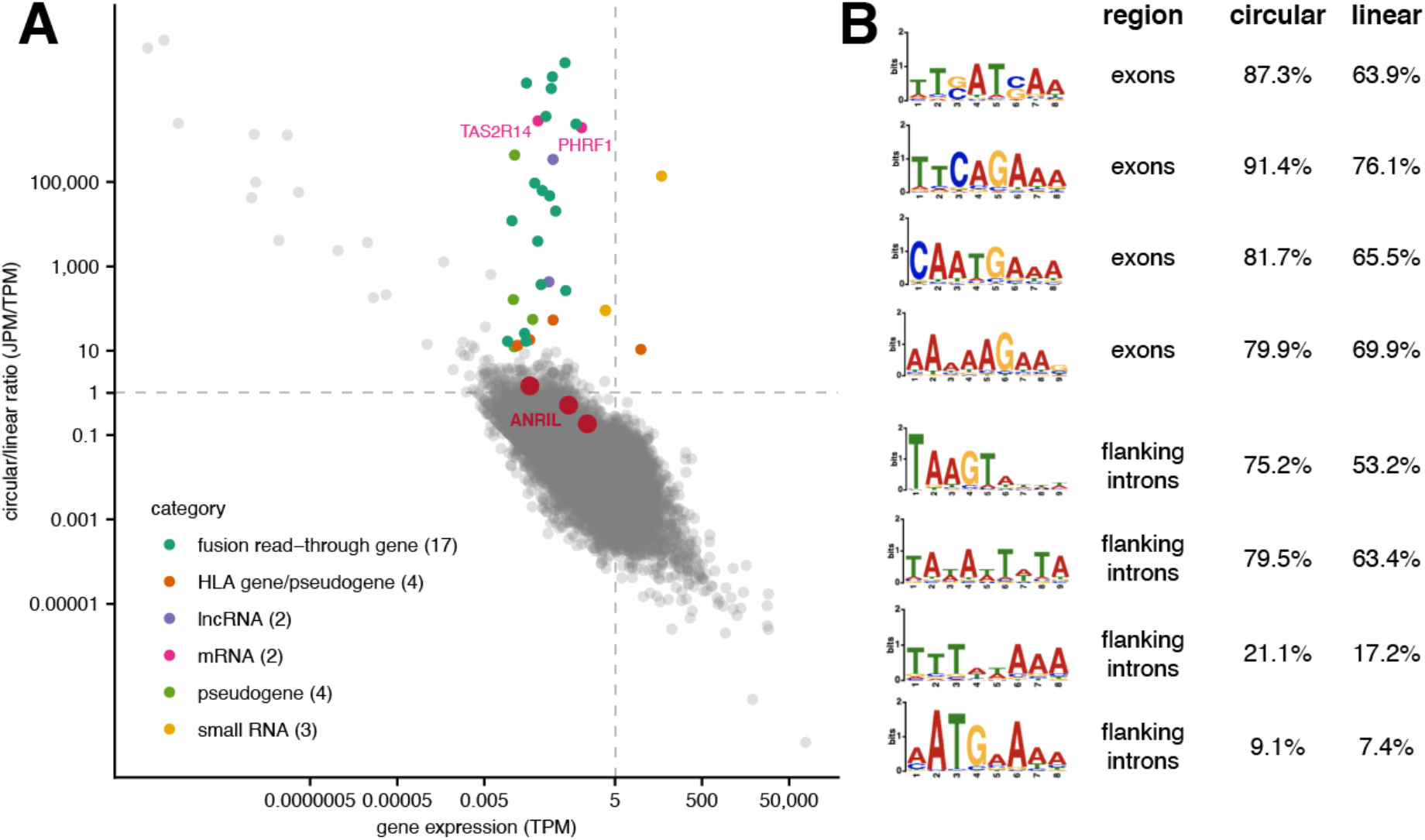
Global expression of circular RNAs in pancreatic islets. **(A)** Correlation between linear gene expression (TPM, *x-axis*) and circular/linear ratios (JPM/TPM, *y-axis*) for all genes with circRNAs in pancreatic islet cells. Genes that diverge from the global correlation patterns are highlighted, with colors differentiating by gene type or functional category. Dotted lines indicate equal circular/linear abundance (*horizontal*) and a threshold for expressed genes (*vertical*). **(B)** Sequence motifs enriched in circRNA-involved exons or flanking introns, with percent occurrence in circular or non-circular (linear) regions indicated on the right.

## DISCUSSION

In recent years, both linear and circular forms of the non-coding RNA ANRIL have been implicated in cardiometabolic disease. While studies have probed the involvement of ANRIL in cardiovascular disease susceptibility and phenotypes (17, 20, 21), less is known about how ANRIL plays a role in T2D, despite multiple strongly validated T2D risk polymorphisms within or near this gene locus (13). Building on our prior observation that T2D risk-SNPs were associated with increased ANRIL abundance and decreased beta cell proliferation index, in this study we used high-throughput sequencing to systematically characterize the identity and abundance of circANRIL isoforms in human pancreatic islet cells. We were able to identify a diversity of high abundance circANRIL isoforms, many of which have been previously identified in other cell types and characterize features that may regulate circularization in islets. Our most important findings include the characterization of features that may regulate ANRIL circularization in islets and association of circ/lin ANRIL ratios with cellular proliferation and an exonic T2D-associated risk SNP.

Previous studies seeking to identify circANRIL molecules have mostly used Sanger sequencing after divergent primer PCR (17, 18, 20, 30, 47), which limits the detection of isoforms due to primer selection and placement. Our unbiased high-throughput sequencing approach allowed us to identify multiple distinct isoforms for circANRIL, with extensive alternative splicing leading to varied internal exon structures. Our results expand the repertoire of isoforms and the exonic or intronic sequences that could be involved in regulation of or by circANRIL. While other studies in immune or cancer cells have mostly focused on one predominant isoform (17, 18, 20), we observe that at least 5 isoforms are relatively abundant in islets. This includes isoforms 7-5, 10-5, and 16-5, which were previously identified and characterized in blood monocytes (17) and T lymphocytes (20) using clonal sequencing. Our observations may reflect cell type heterogeneity, where each individual cell type within islet populations may have a specific predominant isoform, or point to extensive intra-cellular circANRIL diversity that was underappreciated in previous studies.

Across all circANRIL isoforms in islets, a clear pattern emerged that back-splicing mostly involved a specific set of splice sites: 5’ splice sites from exons 7, 10, and 16 and 3’ splice sites from exons 2, 4, and 5. These observations suggest that back-splicing does not occur with equal probability at every ANRIL splice site, but rather, features of these exons or splice sites may promote back-splicing. While we and others (18) see small differences in intron length, sequence complementarity, and motif enrichment for genomic regions flanking these splice sites, there is no one primary differentiating feature. It appears that the frequency of back-splicing is regulated, since circular isoform expression was correlated across individuals. Notably, the abundance of circANRIL isoforms were more correlated with each other than with linANRIL or other mRNAs transcribed from the locus, suggesting independent, non-transcriptional regulation of these distinct ANRIL molecules. Previous studies have also seen that overall circANRIL abundance may be more correlated with CDKN2A and CDKN2B isoforms and perhaps even regulate the abundance of these mRNAs (30). Together, these results highlight the need for a greater understanding of circANRIL regulation across different cell types.

While there are many proposed potential functions for circRNAs, one enduring idea is that circRNAs act as sponges that sequester miRNAs away from their intended targets (79, 80). For this to occur, the circRNA must have target sites for the specific miRNA, with an enrichment of target sites relative to the frequency expected by chance. A previous unbiased bioinformatic analysis trying to identify miRNA sites in circANRIL only identified 3 miRNA target sites in circANRIL (20). However, this study focused solely on isoform 7-5, which only has 3 exons and is expressed almost at the same level as other isoforms containing many more exons in islet cells. Using these longer isoforms, we are able to identify many more miRNA target sites across exons involved in circANRIL and/or linANRIL. Notably, we find that exons involved in circANRIL have a 4-fold enrichment of target sites for miR-130-5p, which has been previously implicated in pancreatic cancers. More work is needed to evaluate the direct binding of the bioinformatically enriched miRNAs we identify and the role of circANRIL-miRNA interactions in islet cells. While previous studies have experimentally characterized miRNA binding to both linear and circular ANRIL molecules in various cell types and cellular contexts (62–65), often the effect is detectable only after cellular stimulation or injury. Similarly, other prominent circRNAs thought to act as miRNA sponges, like CDR1as, have stronger effects after cellular stimulation in islet cells (81). Our analyses were performed on pancreatic islets under basal conditions.

Intriguingly, we find that the ratio of circular to linear ANRIL abundance is associated with beta cell proliferation index and with genotype variation at a T2D-associated SNP. The balance between linear to circular ANRIL species has been previously implicated in regulating the balance between proliferative and apoptotic phenotypes in atherosclerosis models (17, 20). That study proposed a model for cardiovascular disease, in which increased atherosclerosis risk is associated with increased linear ANRIL; linear ANRIL epigenetically regulates factors that promote cell adhesion and proliferation by acting as a molecular scaffold for the polycomb repressive complex (PRC1/2). In contrast, they posit that protection against atherosclerosis is associated with increased circANRIL, where circANRIL protects against over-proliferation by binding and impairing PES1 function to impede ribosomal RNA maturation (20). Similar effects were observed in human brain microvascular endothelial cells, in which circANRIL overexpression inhibited proliferation and apoptotic phenotypes (64) These effects were the most pronounced upon cellular stress. Our observations suggest that the balance between circANRIL and linANRIL may similarly play a role in regulating proliferation of pancreatic beta cells.

Our study has several limitations. Our RNA sequencing and gene expression analyses were performed on whole pancreatic islets, which contain endocrine cells including alpha, beta, delta, and other cells, as well as non-endocrine cells such as endothelial cells. The cell-type distribution of the ANRIL isoforms studied has not been assessed. However, our major results were validated in the endoC-βH1 cell line, an established human beta cell line (40). The insulin secretion analyses included here were based on glucose stimulated insulin secretion testing performed on-site at each human islet isolation center (12); inter-site variation, as well as the high individual variation already noted in human islet preparations (82) may have obscured findings. Furthermore, the majority of samples analyzed here are from non-diabetic individuals. The expression of both linear and circular ANRIL isoforms may be quite dynamic in individuals with T2D or under diabetes-related stresses. It is even possible that different ANRIL isoforms are expressed under these conditions. Finally, despite a combined high-throughput and targeted approach to identify and quantify circANRIL isoforms in islet cells, we were limited by the relatively low abundance of ANRIL RNAs. The need for RNase R to enrich for circRNA and a paucity of BSJ reads across samples hindered our ability to quantify ANRIL isoforms at a high resolution. Techniques to overexpress this locus - either through in-vivo stimulation or experimental perturbation - may prove useful in alleviating this issue.

In sum, our work reveals exciting new biology of the ANRIL lncRNA, including detailed exploration of its circular isoforms in primary human pancreatic islet tissue. Future studies are needed to dissect the regulation of ANRIL circularization, as well as the molecular mechanisms by which linear and circular ANRIL molecules influence beta cell proliferation under basal, stimulated, and stress conditions. Whether ANRIL isoforms impact other important parameters of islet function such as glucose sensing, insulin secretion and cell survival under stress remains unknown. Given the marked worldwide increase in T2D and associated social costs, and the robust relationship between ANRIL locus SNPs and T2D risk, further studies are warranted.

## AVAILABILITY

All software used in this study are from open-source collaborative initiatives or public repositories. Bioinformatics software available in GitHub repositories include Trimmomatic (https://github.com/timflutre/trimmomatic), STAR (https://github.com/alexdobin/STAR), Kallisto (https://github.com/pachterlab/kallisto), CIRCexplorer2 (https://github.com/YangLab/CIRCexplorer2/), CIRIquant (https://github.com/bioinfo-biols/CIRIquant), HTSeq (https://github.com/simon-anders/htseq), and miRanda (https://github.com/hobywan/miranda).

The 5’ splice site and 3’ splice maxEntScan softwares are available at http://hollywood.mit.edu/burgelab/maxent/Xmaxentscan_scoreseq.html and http://hollywood.mit.edu/burgelab/maxent/Xmaxentscan_scoreseq_acc.html, respectively. The BLASTN software to query nucleotide alignments is available through the NCBI website (https://blast.ncbi.nlm.nih.gov/). The STREME program for discovering ungapped motifs is available through the MEME Suite (https://meme-suite.org/meme/doc/streme.html).

## ACCESSION NUMBERS

High-throughput sequencing data have been deposited with the NCBI Gene Expression Omnibus under accession number GSE192541.

## SUPPLEMENTARY DATA

Supplementary Data are available at NAR online.

## ACKNOWLEDGEMENT

We thank members of the Alonso and Pai labs for discussion and comments on the manuscript, as well as the anonymous donors who contributed their pancreatic islet cells to medical research. Some pancreatic islets were graciously provided by Drs. Alvin Powers and Marcela Brissova at Vanderbilt University Medical Center, who organized the collection of de-identified human pancreatic islet samples as part of NIH-funded grants (DK106755, DK104211, and DK108120).

## FUNDING

This work was supported by American Diabetes Association in collaboration with the Order of the Amaranth [1-18-IBS-233] to LCA and National Institute of General Medical Sciences [R35GM133762] to AAP. Human pancreatic islets were provided by the Integrated Islet Distribution Program (IIDP) funded by National Institute of Diabetes and Digestive and Kidney Disease [RRID:SCR _014387, 2UC4DK098085] to City of Hope. Funding for open access charge: National Institutes of Health R35GM133762.

## CONFLICT OF INTEREST

None declared.

## Notes

### Competing Interest Statement

The authors have declared no competing interest.

