## Supplementary Figures for "High-throughput analysis of ANRIL circRNA isoforms in human pancreatic islets"

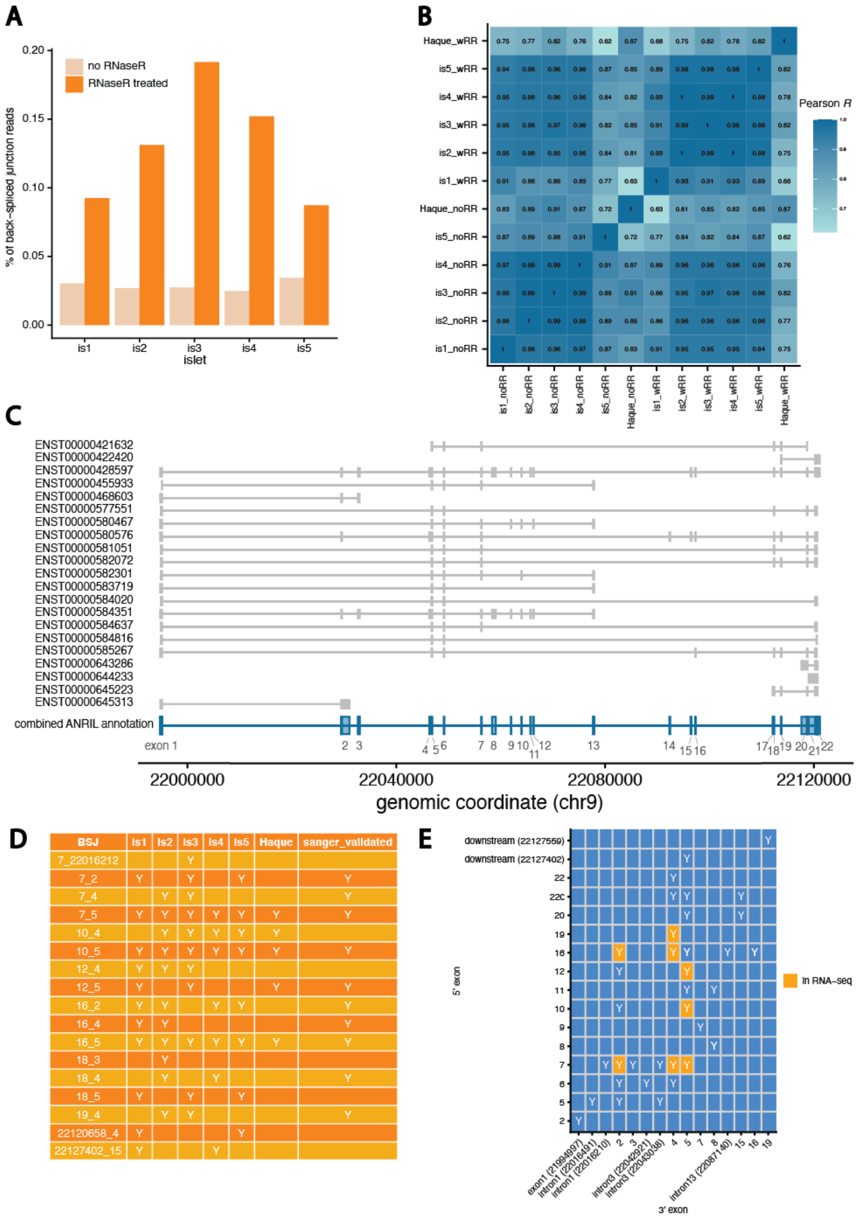


**Supplementary Figure 1. Identifying circular ANRIL isoforms. (A)** Percentage of reads that represent back-spliced junctions across control (*beige*) and RNaseR treated (*orange*) samples from each islet preparation. **(B)** Pairwise Pearson correlations of gene expression levels across samples from 5 islet preparations and published data from Haque *et al*. **(C)** Annotated ANRIL isoforms that were combined to create a meta-isoform (*blue*). **(D)** Back-spliced junction positions identified from RNaseR RNA-seq samples across each islet preparation and Haque *et al.* data, after filtering for read count and minimum number of samples. **(E)** Back-spliced junction positions identified in Sanger sequencing data. Junctions concordant with RNA-seq data are in orange.

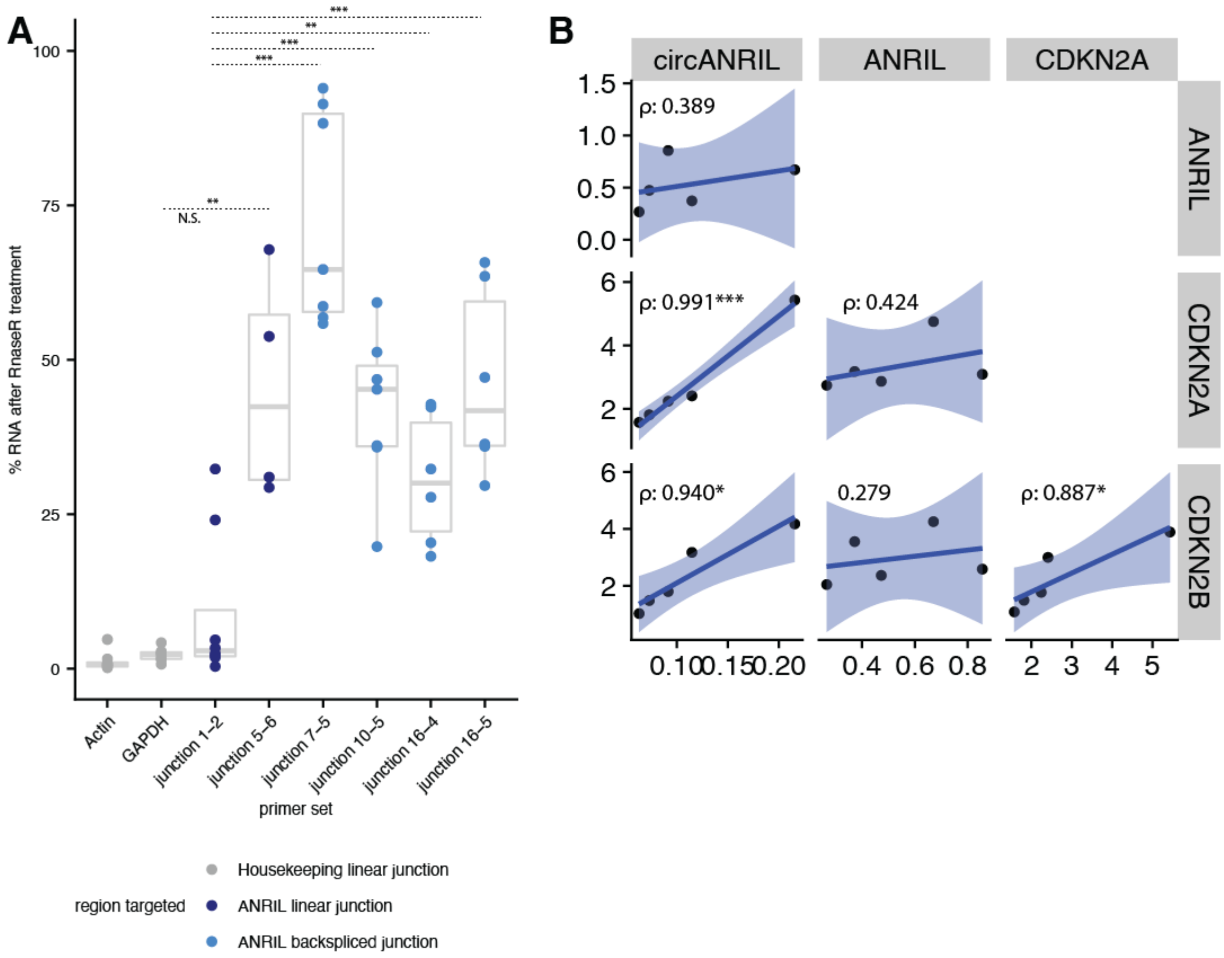


**Supplementary Figure 2. Quantification of ANRIL isoforms.** (A) RT-PCR quantification of linear and circular ANRIL isoforms in endoC-βH1 cells (** p-value < 0.01, *** p-value < 0.001). (B) Pairwise Spearman correlations between circular ANRIL (JPM), linear ANRIL, CDKN2A, and CDKN2B (all TPM) RNA-seq derived expression levels across 5 islet preparations.

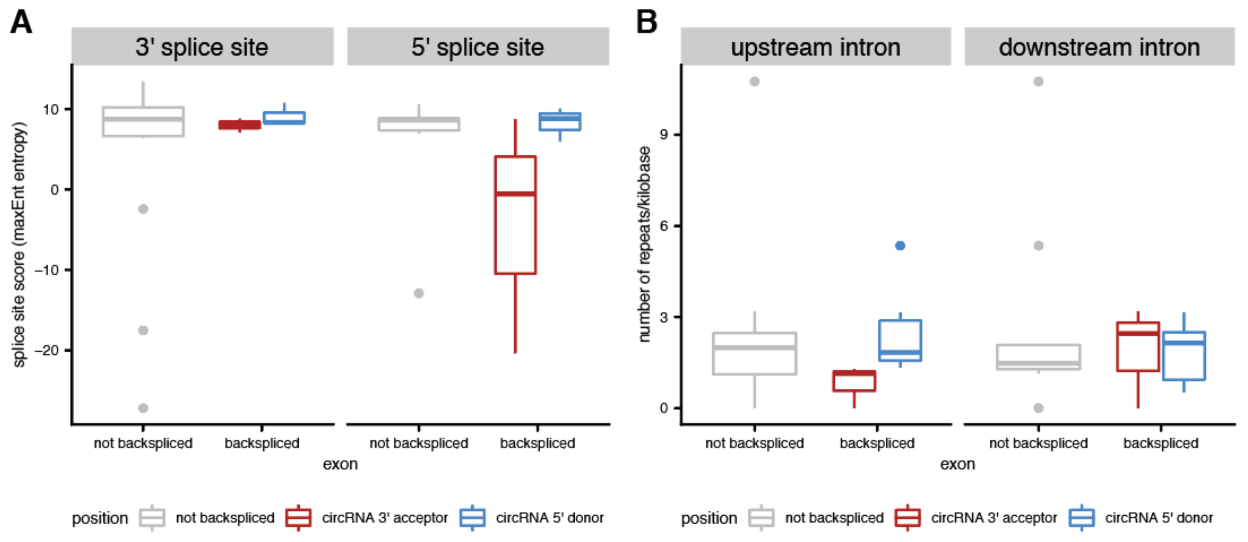


**C**

| Foreground | Background | Sequence | Train (Fg) | Train (Bg) | Score | % Match (Fg) | % Match (Bg) | Sites | FC (Fg/Bg) | RBPdb Hit |
|---|---|---|---|---|---|---|---|---|---|---|
| BSJ involved exons | non−BSJ involved exons | AACACTGGG | 9 of 11 | 0 of 11 | 0.00011 | 81.80% | 0.00% | 9 | 81801.000000 | SFRS13A |
| BSJ involved 5' ss introns | non−BSJ involved 5' ss introns | TAAATAAAA | 10 of 10 | 2 of 11 | 0.00019 | 100% | 18% | 10 | 5.555302 | KHDRBS3 |
| BSJ involved 5' ss introns | non−BSJ involved 5' ss introns | AGATACTC | 9 of 10 | 1 of 11 | 0.00031 | 90% | 9% | 9 | 9.999000 | NA |
| BSJ involved 5' ss introns | non−BSJ involved 5' ss introns | ATWATAGTTM | 9 of 10 | 1 of 11 | 0.00031 | 90% | 9% | 9 | 9.999000 | NA |
| BSJ involved 5' ss introns | non−BSJ involved 5' ss introns | CTCTCTTCA | 10 of 10 | 3 of 11 | 0.00081 | 100% | 27% | 10 | 3.703604 | NA |

**Figure S3. Regulation of ANRIL circRNAs. (A)** 3' and 5' splice site maxEnt scores (*y-axis*) for BSJ-involved exons and exons not involved in BSJs. **(B)** Distribution of the number of known repeat regions (*y-axis*) in introns located upstream (*left*) and downstream (*right*) of exons involved in BSJs or not involved in BSJs. **(C)** Table of motif enrichments output by STREME showing enriched motifs information and putative RBPs matches from RBPDB. FC indicates fold change between foreground (BSJ-associated) and background (non-BSJ-associated) sequences.

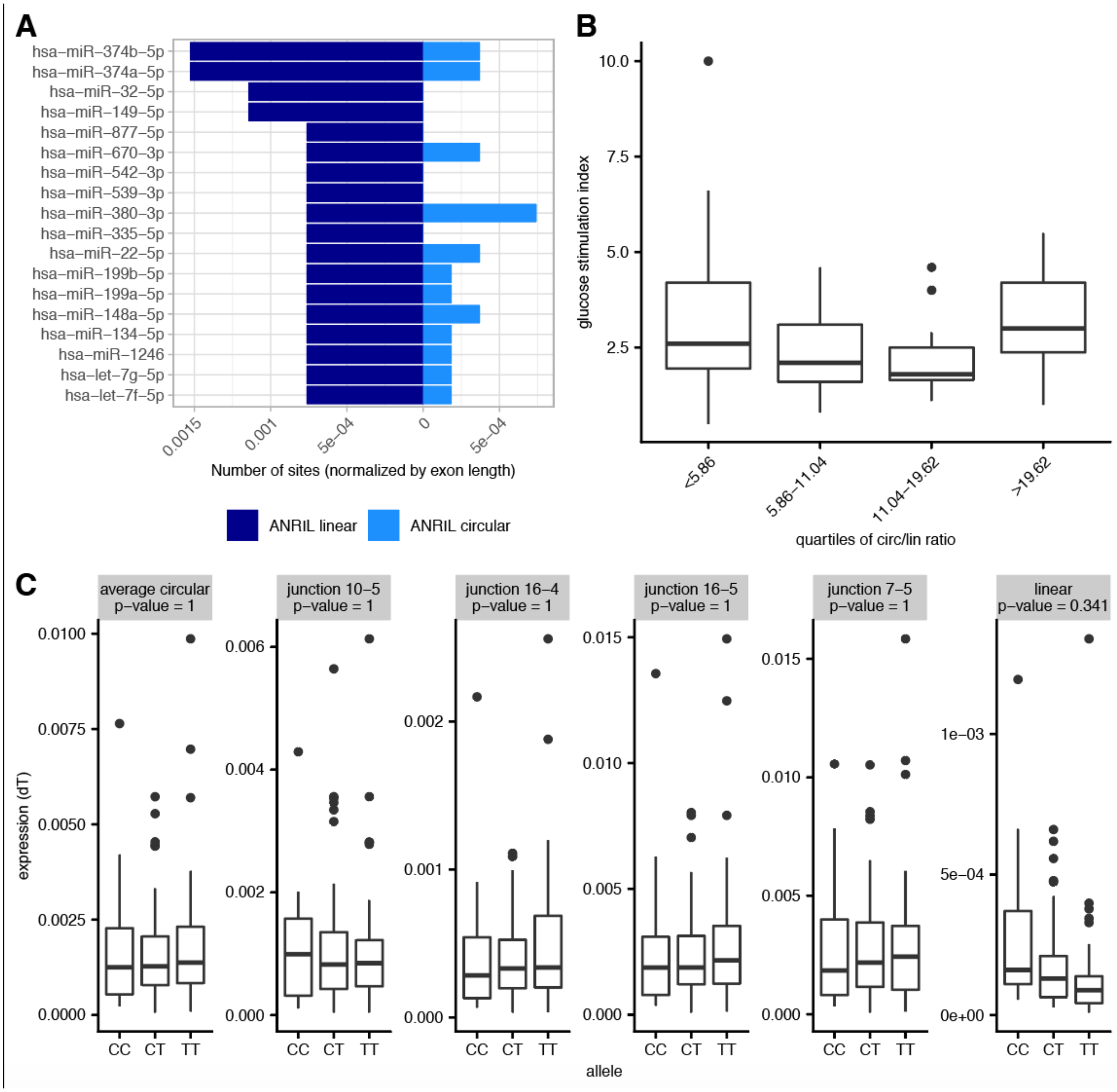


**Figure S4: circANRIL functional associations. (A)** Number of miRNA target sites (normalized by exon length, *x-axis*) in ANRIL circular exons (*light blue*) vs ANRIL linear exons (*dark blue*) for miRNAs with more sites in linANRIL exons. **(B)** Distribution of glucose stimulation indices (*y-axis*) for quartiles of circANRIL/linANRIL ratios (*x-axis*) as measured by RT-PCR across 83 islet preparations. **(C)** Association between the genotypes of rs564398 and ANRIL isoform expression across 122 islet preparations. P-values are computed from a linear regression and Benjamini-Hochberg corrected.

A

| Foreground | Background | Sequence | Train (Fg) | Train (Bg) | Score | % Match (Fg) | % Match (Bg) | Sites | FC (Fg/Bg) | RBPdb Hit |
|---|---|---|---|---|---|---|---|---|---|---|
| BSJ involved exons | non–BSJ involved exons | ATGAAGAA | 13803 of 14964 | 6323 of 8670 | 3.3E–341 | 0.922 | 0.729 | 15310 | 1.26 | NA |
| BSJ involved exons | non–BSJ involved exons | TTCTGAA | 12468 of 14964 | 5570 of 8670 | 7.2E–236 | 0.833 | 0.642 | 13807 | 1.30 | NA |
| BSJ involved exons | non–BSJ involved exons | ATCAWTGAT | 13140 of 14964 | 6523 of 8670 | 9.6E–133 | 0.878 | 0.752 | 14592 | 1.17 | NA |
| BSJ involved exons | non–BSJ involved exons | YKMCAAYN | 13908 of 14964 | 7235 of 8670 | 1.1E–111 | 0.929 | 0.834 | 15416 | 1.11 | NA |
| Exons in genes with circRNAs | Exons in genes without circRNAs | TAATGAAA | 1073 of 10183 | 635 of 10545 | 1.2E–26 | 0.105 | 0.060 | 1189 | 1.74 | NA |
| Exons in genes with circRNAs | Exons in genes without circRNAs | AATATGTA | 624 of 10183 | 309 of 10545 | 1.7E–25 | 0.061 | 0.029 | 691 | 2.07 | NA |
| Exons in genes with circRNAs | Exons in genes without circRNAs | AAATTATT | 3966 of 10183 | 2917 of 10545 | 7.5E–37 | 0.389 | 0.277 | 4405 | 1.40 | NA |
| Exons in genes with circRNAs | Exons in genes without circRNAs | AAAAGCTTTAA | 537 of 10183 | 260 of 10545 | 3E–23 | 0.053 | 0.025 | 595 | 2.08 | NA |
| BSJ involved 5'ss | non–BSJ involved 5'ss | TAAGTAWWW | 27085 of 36000 | 19146 of 36000 | 4.5E–840 | 0.752 | 0.532 | 30109 | 1.41 | ELAVL1 |
| BSJ involved 5'ss | non–BSJ involved 5'ss | TAWAWTWTA | 28627 of 36000 | 22837 of 36000 | 1.0E–503 | 0.795 | 0.634 | 31831 | 1.25 | ELAVL1 |
| BSJ involved 5'ss | non–BSJ involved 5'ss | TTTWWAAA | 7580 of 36000 | 6188 of 36000 | 4.9E–40 | 0.211 | 0.172 | 8386 | 1.23 | KHDRBS3 |
| BSJ involved 5'ss | non–BSJ involved 5'ss | AATGAAAA | 3273 of 36000 | 2649 of 36000 | 1.4E–17 | 0.091 | 0.074 | 3631 | 1.23 | NA |
| BSJ involved 3'ss | non–BSJ involved 3'ss | AAAAGAAAAAAA | 31121 of 36000 | 21662 of 36000 | 3.3E–1429 | 0.864 | 0.602 | 34575 | 1.43 | PABPC1,SFRS13A |
| BSJ involved 3'ss | non–BSJ involved 3'ss | WTAWWWTAW | 25220 of 36000 | 18790 of 36000 | 5.1E–531 | 0.701 | 0.522 | 28041 | 1.34 | NA |
| BSJ involved 3'ss | non–BSJ involved 3'ss | TGTTAATA | 4967 of 36000 | 4322 of 36000 | 4E–13 | 0.138 | 0.120 | 5534 | 1.15 | KHDRBS3 |
| BSJ involved 3'ss | non–BSJ involved 3'ss | TAAAAAAA | 7617 of 36000 | 5875 of 36000 | 1.6E–62 | 0.212 | 0.163 | 8428 | 1.30 | PABPC1 |

**Figure S5. Global circRNA motif analysis.** Table of motif enrichments output by STREME showing enriched motifs information and putative RBPs matches from RBPDB. FC indicates fold change between foreground (BSJ-associated) and background (non-BSJ-associated) sequences.

**Supplementary Table Legends**

**Table S1. Donor information for human islets used for RNA-seq.** Data provided by the Integrated Islet Distribution Program. BMI, body mass index. WIT, warm ischemia time. CIT, cold ischemia time. Initial islet index refers to insulin secretion index measured at the islet isolation center.

**Table S2. ANRIL BSJs identified from RNA-seq data.** Outputs from CIRCexplorer2 (sheet 1) and CIRIquant (sheet 2) for back-spliced junctions in RNaseR treated and untreated samples.

**Table S3.** Concordance of ANRIL back-spliced junction positions identified from RNaseR RNA-seq samples across each islet preparation and Haque et al. data, including all positions before filtering for high-confidence sites.

**Table S4. Sanger sequencing data.** Divergent primers used for Sanger sequencing (sheet 1) and sequencing results for each primer set (sheets 2 - 9), listing genomic coordinates for all non-contiguous regions found in individual clones.

**Table S5. Donor information for human islets used for Taqman qRT-PCR assays.** Data provided by the Integrated Islet Distribution Program. BMI, body mass index. WIT, warm ischemia time. CIT, cold ischemia time. Initial islet index refers to insulin secretion index measured at the islet isolation center.

**Table S6.** Target sites identified in ANRIL exons for miRNAs expressed in human islet cells.

**Table S7. Global BSJs identified from RNA-seq data.** Outputs from CIRCexplorer2 (sheet 1) and CIRIquant (sheet 2) for back-spliced junctions in RNaseR treated and untreated samples.

**Table S8. Quantification of gene-specific circRNA abundance.** CircRNA abundance (in junctions per million) and linear RNA abundance (in transcript per million) for all genes with circRNAs expressed in human islet cells.
